# pH and Receptor Induced Conformational Changes-Implications Towards S1 Dissociation of SARS-CoV2 Spike Glycoprotein

**DOI:** 10.1101/2020.12.21.410357

**Authors:** Jesu E. Castin, Daniel A. Gideon, Karthik S. Sudarsha, Sherlin A. Rosita

**Affiliations:** Department of Biotechnology and Bioinformatics, Bishop Heber College (Autonomous), Tiruchirappalli, India

## Abstract

Viruses, being obligate intracellular parasites, must first attach themselves and gain entry into host cells. Viral fusion machinery is the central player in the viral attachment process in almost every viral disease. Viruses have incorporated an array of efficient fusion proteins on their surfaces to bind efficiently to host cell receptors. They make use of the host proteolytic enzymes to rearrange their surface protein(s) into the form which facilitates their binding to host-cell membrane proteins and subsequently, fusion. This stage of viral entry is very critical and has many therapeutic implications. The current global pandemic of COVID-19 has sparked severe health crisis and economic shutdowns. SARS-CoV2, the etiological agent of the disease has led to millions of deaths and brought the scientific community together in an attempt to understand the mechanisms of SARS-CoV2 pathogenesis and mortality. Like other viral fusion machinery, CoV2 spike (S) glycoprotein- ‘The Demogorgon’ poses the same questions about viral-host cell fusion. The intermediate stages of S protein-mediated viral fusion are unclear owing to the lack of structural insights and concrete biochemical evidence. The mechanism of conformational transition is still unclear. S protein binding and fusion with host cell receptors, Eg., angiotensin-converting enzyme-2 (ACE2) is accompanied by cleavage of S1/S2 subunits. To track the key events of viral-host cell fusion, we have identified (*in silico*) that low pH-induced conformational change and ACE-2 binding events promote S1 dissociation. Deciphering key mechanistic insights of SARS-CoV2 fusion will further our understanding of other class-I fusion proteins

## Background

Viruses constitute a vast majority of space under the infectious disease category. In the history of pandemics, viruses along with bacteria have conferred a major threat to world health as well as the economy. Unlike bacteria, they have higher mutation rates which make them easy to evolve into a better construct. HIV-I genome showed the highest mutation rate of (4.1 ± 1.7) × 10-3 per base per cell **[1]**. The infectivity and lethality can vary from benign to acute. Although there are specific characteristics of each virus that set each of them apart are being crudely focused, there is some common key machinery that helps them evade the host cellular environment. One of such machinery is viral-host membrane fusion mediated by viral glycoproteins and host proteolytic cleavage. Viral glycoproteins also known as fusion proteins, are extremely capable of transitioning into different structurally distinct prefusion/post-fusion states **[2–4]**. Based on these common features of transitions, fusion proteins are classified into three classes. Despite the knowledge about two states, the intermediate events for any viral fusion are poorly understood **[5]**. The novel construct of Coronavirus genome- SARS-CoV2 poses the same inadequacy of knowledge about fusion machinery. Like other fusion proteins, it undergoes spontaneous transition driven by factors like i) low pH **[6]**, ii) Host Receptor Binding **[7]**, iii) Host-mediated proteolytic cleavage **[8]** and iv) host membrane composition (In case of Ebola) **[9]**. Although there are many biochemical analyses **[10–14]** for the intermediates, many structural insights still remain unclear. Spike glycoprotein of SARS-CoV2 is the class I fusion protein with trimeric assembly **[15]**. Deciphering the intermediate stages of the pre-fusion to post-fusion conformational transition can change the whole knowledge about viral fusion proteins. This could lead us to develop effective therapeutics which can block the transition as well as the successful fusion.

## Introduction

The spike glycoprotein of SARS-CoV2 is the large trimeric structure crucial for viral-cell fusion with the molecular weight of about 424 kDa. The structure has the three major topological domains which include Head, Stalk and Cytoplasmic Tail (CT). Each monomeric unit of the protein has two subunits namely S1 and S2 **[16]**. S1 has the regions responsible for host receptor binding-N-terminal Domain (NTD) and Receptor Binding domain (RBD). Most of the S2 regions serve as the fusion machinery-Fusion Peptide (FP), Central Helix (CH), Heptad Repeat (HR), etc (Figure 1.1, Figure 1.2.A). Although these regions are mostly conserved (compared to older strains of Coronaviridae family), few mutations are seen in the Receptor Binding Domain (RBD) and it is also noteworthy that this new construct has polybasic cleavage site **[18]** which is not present in the previously encountered viral strains. This new region increases the rate of S1/S2 cleavage and thereby, elevates efficiency of viral entry into the host cells. Only for the Head domain, the experimental structure is available and the rest of the topological domains have been successfully modelled in accordance with many experimental data **[19]**. The stalk domain is responsible for global protein flexibility and allows the glycoprotein to bend in different angles **[20]**. It is the best example of a mechanistic protein *i.e*., it has exhibited many states which are shown in the Figure1.2 B.

**Figure1.1.**
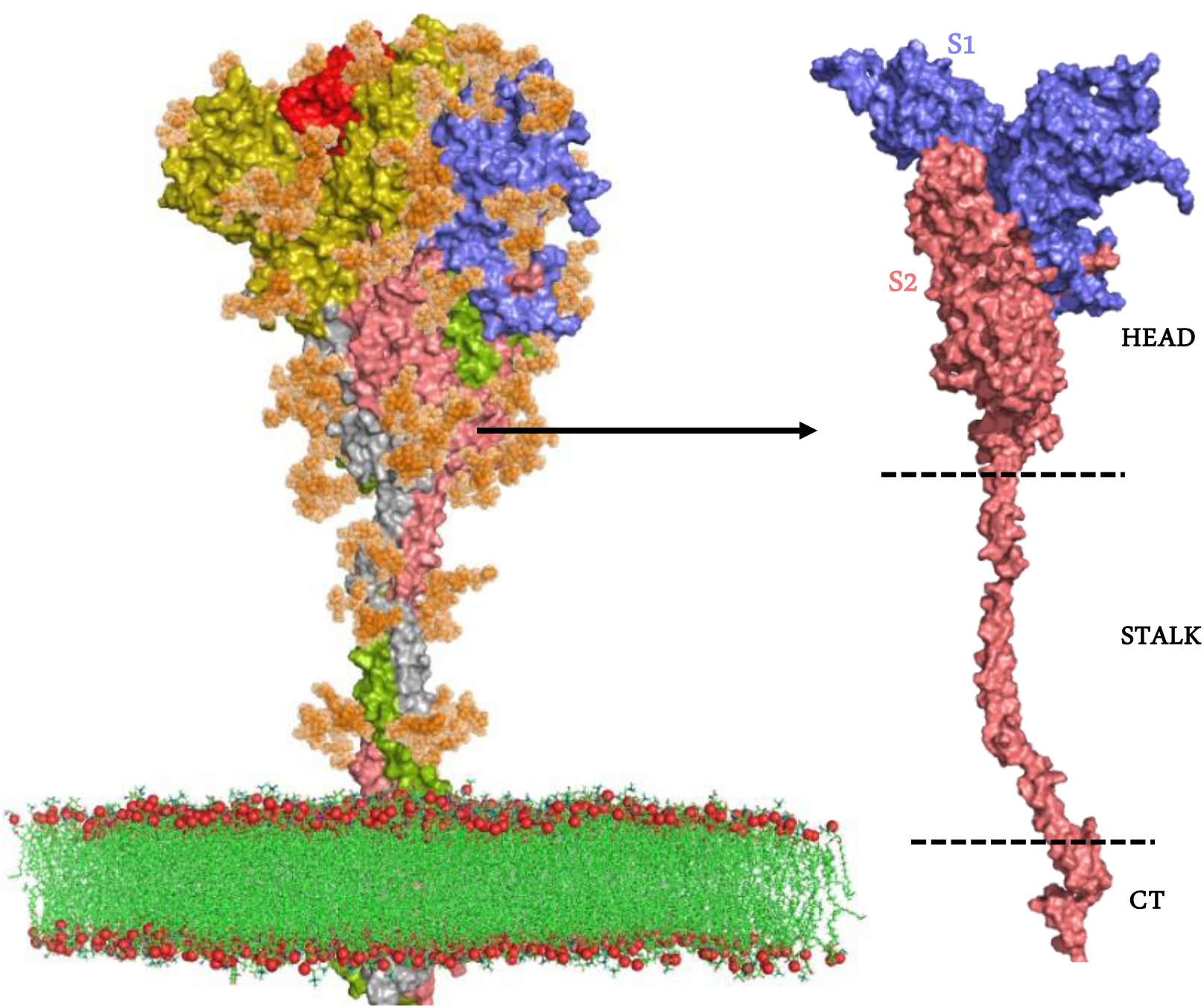
Full Length Glycosylated Spike Model. SARS-CoV2 spike glycoprotein forms a trimeric assembly in which each monomeric unit has two subunits S1 and S2. The main topological domains include head, stalk and cytoplasmic tail (CT).

**Figure 1.2.**
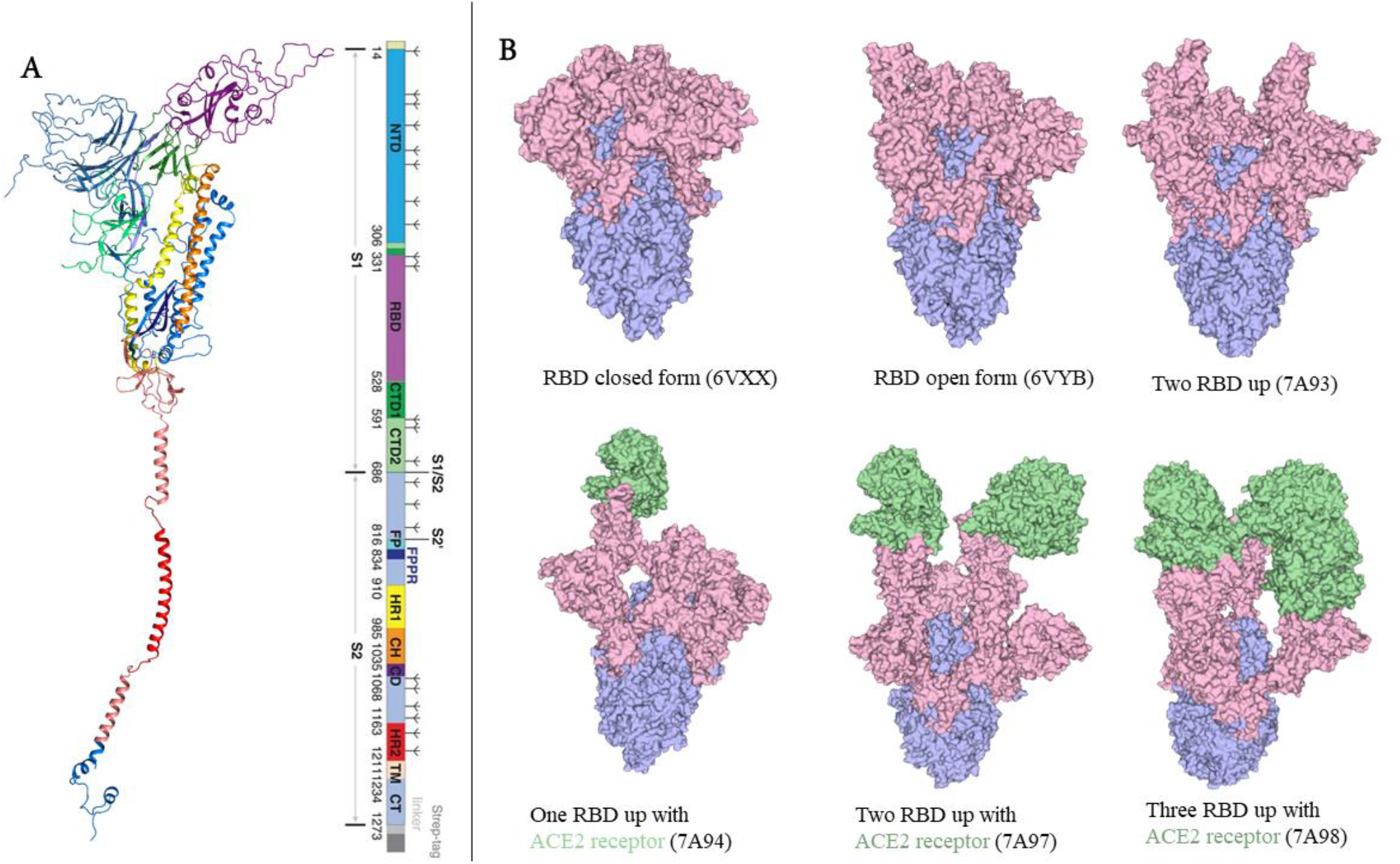
A- Characteristic regions of the spike glycoprotein (Chain A). Spike has several key regions which are responsible for various functions that promote the viral entry into the host cell. Each region is coloured with respective to the one-dimensional legend. **Figure 1.2 B- Spike Exhibiting Multiple States (with the PDB ID)**

Followed by Angiotensin Collecting Enzyme-2 (ACE-2) Receptor binding in open form/ RBD ‘Up’ form (active form), the fusion events are initiated. The cleavage is likely to be mediated by host furin or TRPMSS2 at two distinct sites-S1/S2 and S2/S2’ respectively **[21]**. As stated earlier, the polybasic cleavage site at the S1/S2 interface brings about effective proteolytic cleavage, thereby promoting the mechanistic tendency of the spike glycoprotein to mature into the post-fusion state. The steps of such transitions are unclear because very few biochemical evidence has been found for the intermediates.

But it is well known that S1/S2 cleavage (Figure 1.3. A) promotes S1 dissociation whereas S2/S2’ (Figure 1.3. B) cleavage paves the way for S2 maturation (into the postfusion state) because Fusion peptide carrying the membrane fusing segment- ‘FPPR’ is released along with Heptad Repeat (HR) which holds the FP straight towards the host membrane. Although the cleaved states for the Stabilized CoV1 did not show any immediate conformational changes in the Cryo-EM experiments **[22]**, our analysis on CoV2 Spike showed that the post cleavage state is much more disordered. Right after the binding event with ACE-2, the structural changes like decreased S1/S2 contact start to occur **[23]**. Our preliminary analysis confirmed this conformational change and yielded deeper insights.

**Figure 1.3-.**
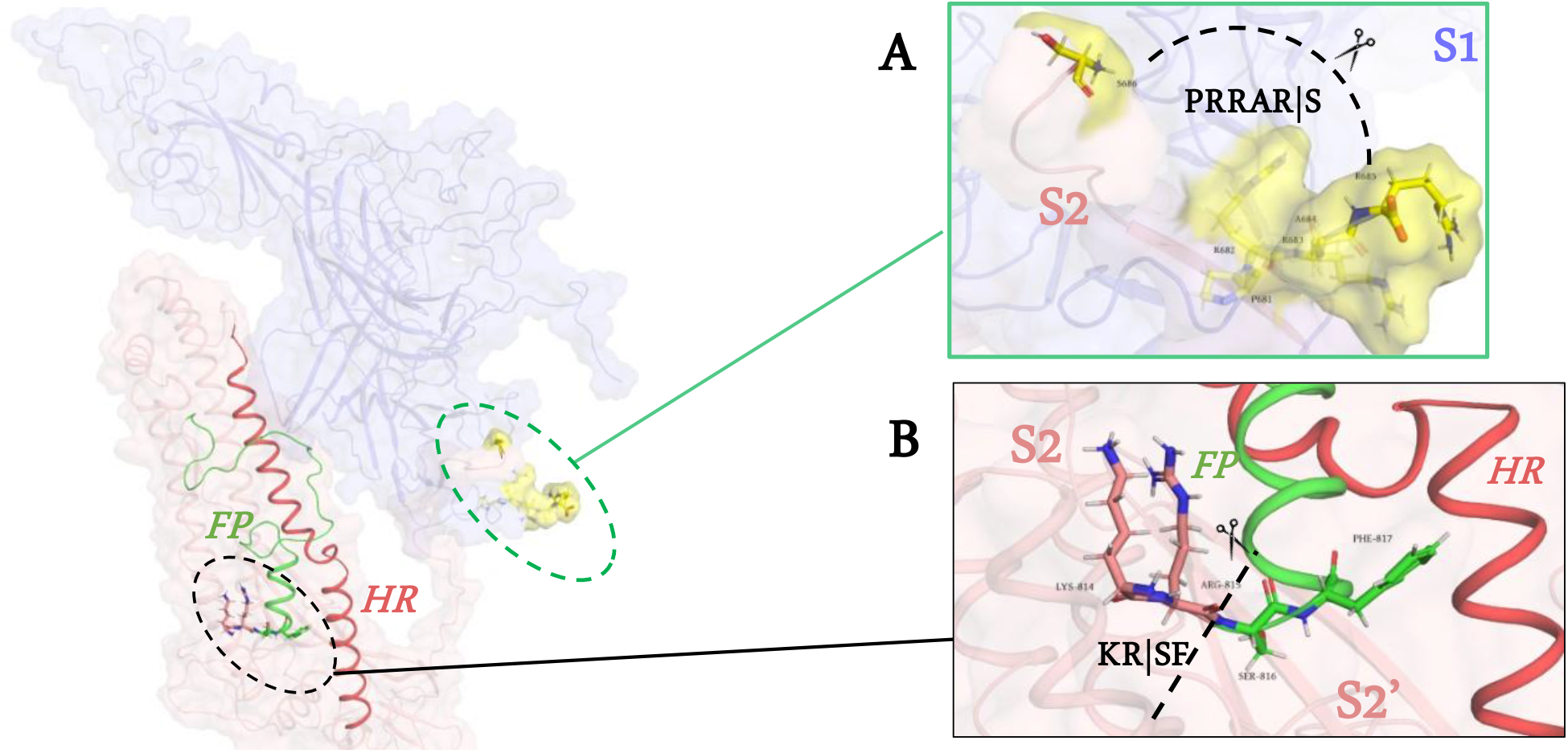
S1/S2 and S2/S2’ cleavage sites

The key role of glycans in protein shielding has come to the fore recently. Due to the recent reports, several glycan associated analyses have become prominent and repositories (such as protein data bank) have focused on easier retrieval of glycans which are associated with proteins. A recent study on SARS-CoV2 spike by Casalino et al., 2020 has shed more light on the role of glycans beyond just protein shielding. In that study, the group identified that the glycans in N165 and N234 stabilize the RBD ‘Up’ state **[24]**. And another study depicted its importance in maintaining contact with the ACE-2 Receptor **[25]**. Recently, the same glycans involved in RBD ‘Up’ state are known to form a cryptic pocket during the intermediate’s opening pathway **[26]**.

## Results and Discussion

### i) Low pH creates unfavourable charge distribution in S1/S2 Interface and thereby promotes S1 Dissociation

S1 dissociation is a spontaneous step which occurs after S1/S2 Cleavage and the ACE-2 Binding event. S1 and S2 have two putative contact interfaces C1 and C2 (Supplementary Figure 2.1). Of the two, Amaro Lab ‘Closed’ state model (depicting postcleaved S1/S2 state) showed that C2 has a complex hydrogen-bonding network (Supplementary Figure 2.1). Also, a strong electrostatic complementarity was seen in the C1 and C2 (Supplementary Figure 2.2). But in the dynamical events, the external physical conditions were found to play a vital role in the conformational arrangement of proteins. Hence, we hypothesised that the low pH would create an unfavourable charge distribution. The hypothesis is successfully proven by the preliminary results from ProPka. The pKa values for the titratable residues are calculated as follows:

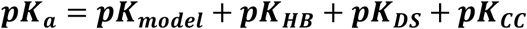

where ***pK_model_*** is the reference pKa value for each titratable amino acid and ***pK_HB_, pK_DS_, pK_CC_*** are the pKa values decreased in the event of hydrogen bonding interaction, desolvation and coulombic contact, respectively.

On analysing the titratable residues in the S1/S2 Interface (Figure 1.4.A) of ‘Closed’ and ‘ACE-2 Bound’ experimental structures (PDB ID: 6VXX and 6A98 respectively), it becomes clear that the interface residues of the ‘ACE-2 Bound’ state exhibited higher pKa values than the interface residues of the ‘Closed’ state (Figure 1.4.B). This implies that the hydrogen bonding network and Coulombic contacts begin to diminish at the low pH receptor binding event, which would result in increased ***pK_HB_, pK_CC_*** values. Also, charge calculations across the pH spectrum proved that low pH induces unfavourable positive/neutral charge distribution in the S1/S2 interface (Figure 1.5).

**Figure 1.4.**
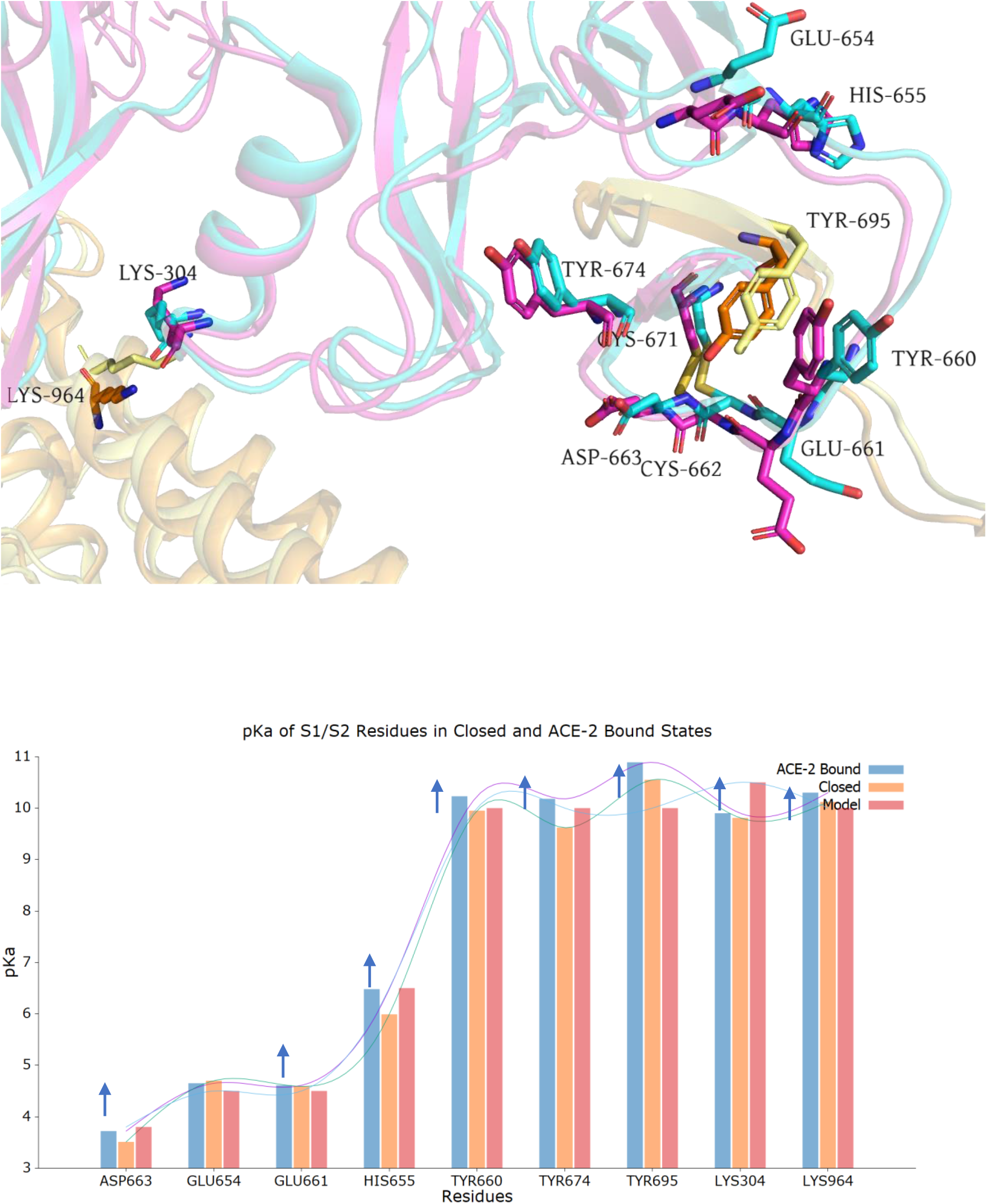
A- Titratable residues at the S1/S2 interface of Closed (PDB: 6VXX) and ACE-2 Bound states (PDB: 6A98) **Figure 1.4.B- pKa values of S1/S2 interface titratable residues**. The pKa values of the interface residues in ‘ACE-2 Bound’ state (shown in arrows) are increased due to the diminishing polar and coulombic contact

**Figure 1.5-.**
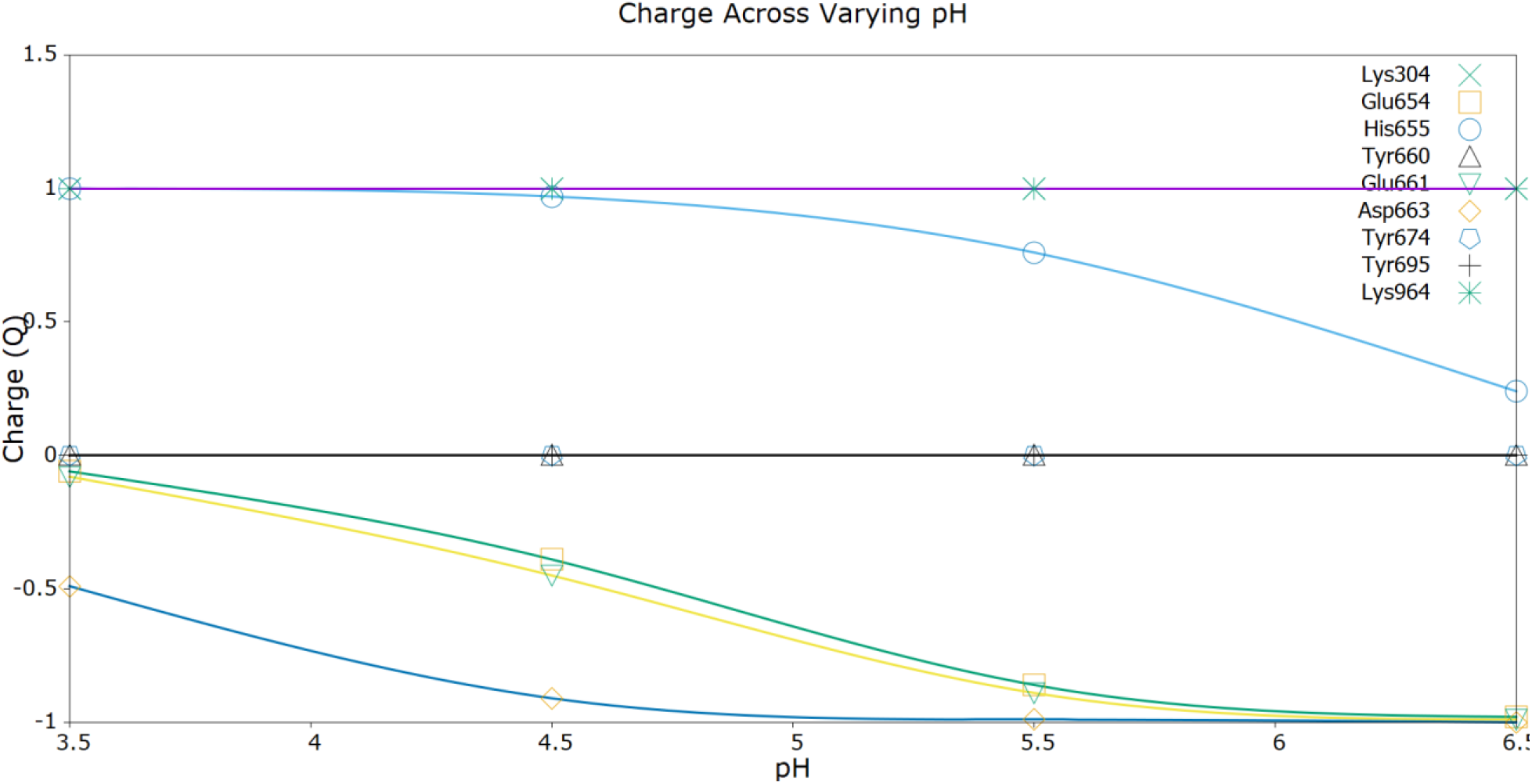
Charge calculation across the decreasing pH. The graph implies that most of the interface residues show increasing positive/neutral charge distribution when the pH decreases (other than lysine and tyrosine residues)

We obtained a different result from Amaro lab model of ‘Open’ state (Supplementary figure 2.4.A). Unlike in ‘ACE-2 Bound’ state, the pKa values of the interface residues are decreased, providing us with the insights on the differences between ‘Open’ and ‘ACE-2 Bound’ states. The charge distribution again showed increasing values across decreasing pH (Supplementary figure 2.4.B).

### ii) S1/S2 Cleavage Elevates Local Disorderliness in S1/S2 Interface Environment

We compared the uncleaved (precleaved) (CHARMM Model) and postcleaved (Amaro Lab Model) S1/S2 models (Figure 1.6) with Normal Mode Analysis incorporated with Anisotropic Network Model (ANM). NMA is the approximation technique of Molecular Dynamic Simulation that uses a network model and calculates projection of each atom assuming a harmonic oscillation. Calculating Hessian Matrix for each atomic projection can provide us with insights about fluctuations and other dynamical properties. Applying NMA-ANM for the postcleaved as well as precleaved states, we came to find that postcleaved S1/S2 state showed increased RMSF (especially in S1/S2 interface) than that of precleaved S1/S2 state (Figure 1.7). It provided the preliminary insight that host-mediated proteolytic cleavage elevates relative disorderliness in S1/S2 interface and further promotes S1 dissociation. It is noteworthy that neither experimental structures nor models are capable of illustrating S2/S2’ cleavage, which can be studied only through rigorous MD approaches.

**Figure 1.6-.**
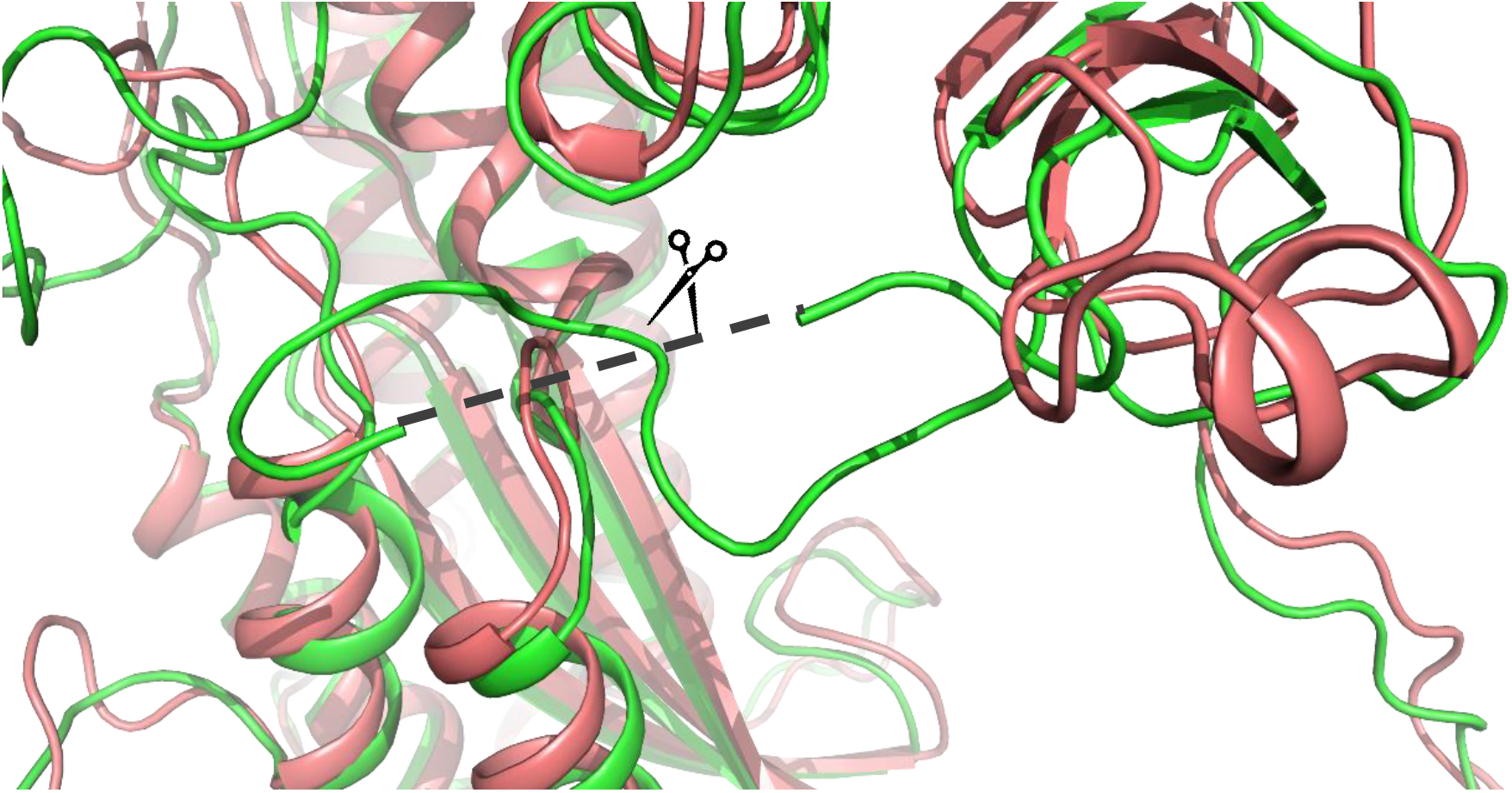
Superimposed precleaved (CHARMM Model) and postcleaved (Amaro Lab Model) states.

**Figure 1.7-.**
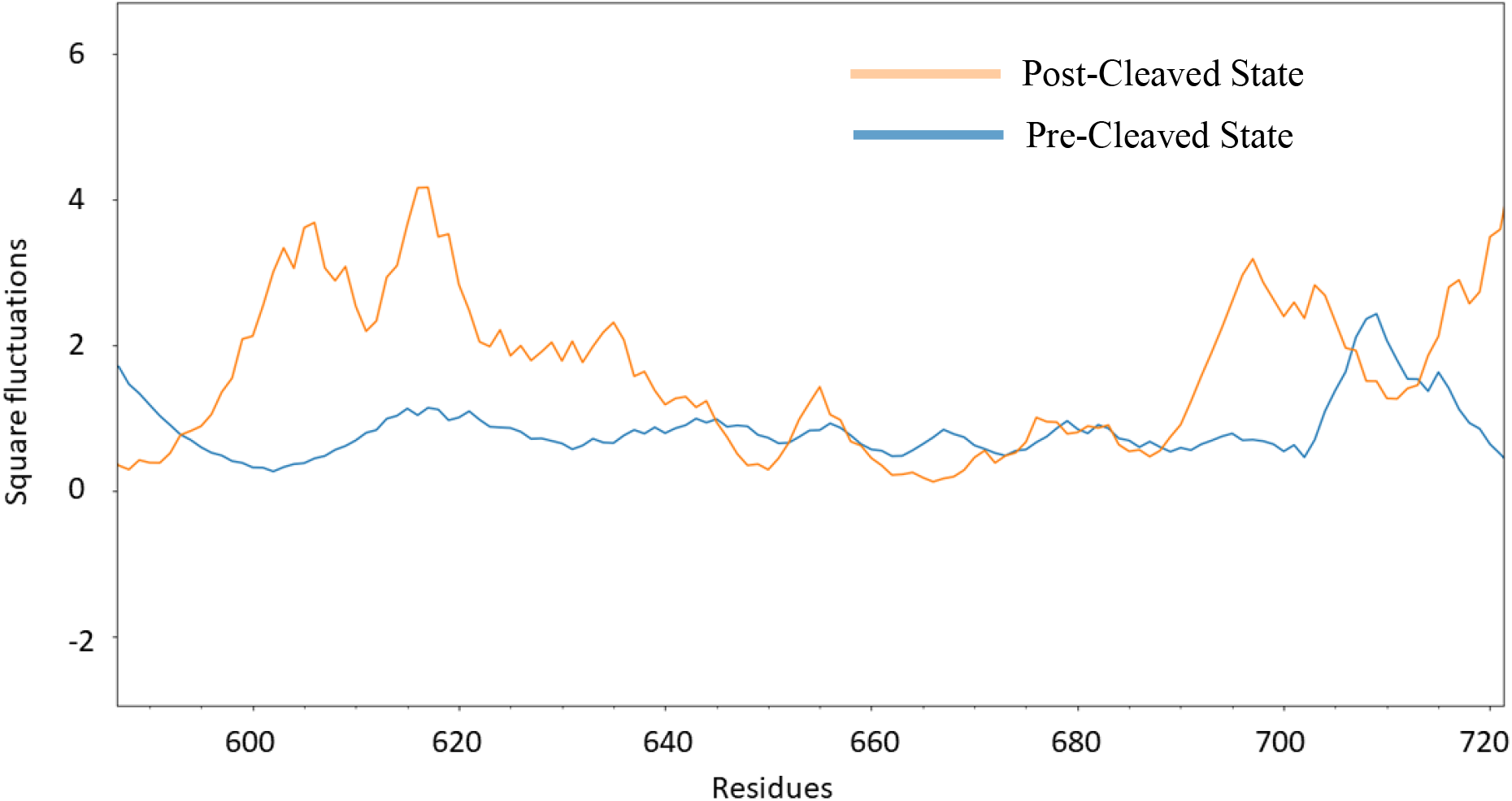
RMSF plot shows that the S1/S2 interface in post-cleaved state is highly fluctuating in nature.

### iii) Experimental Evidence Proves Host-Receptor Induced Conformational Changes

We also generated distant maps for the experimental structures in ‘Closed’ and ‘ACE-2 Bound’ states (Figure 1.8). It became clear that the contact distance in S1/S2 interface of the ‘ACE-2 Bound’ state appeared to have diminished. We could observe that the NTD and CTD-2 regions are fluctuating in nature whereas HR of S2 remained stable (Figure 1.9). This confirms that Host-Receptor binding also influences S1 dissociation.

**Figure 1.8-.**
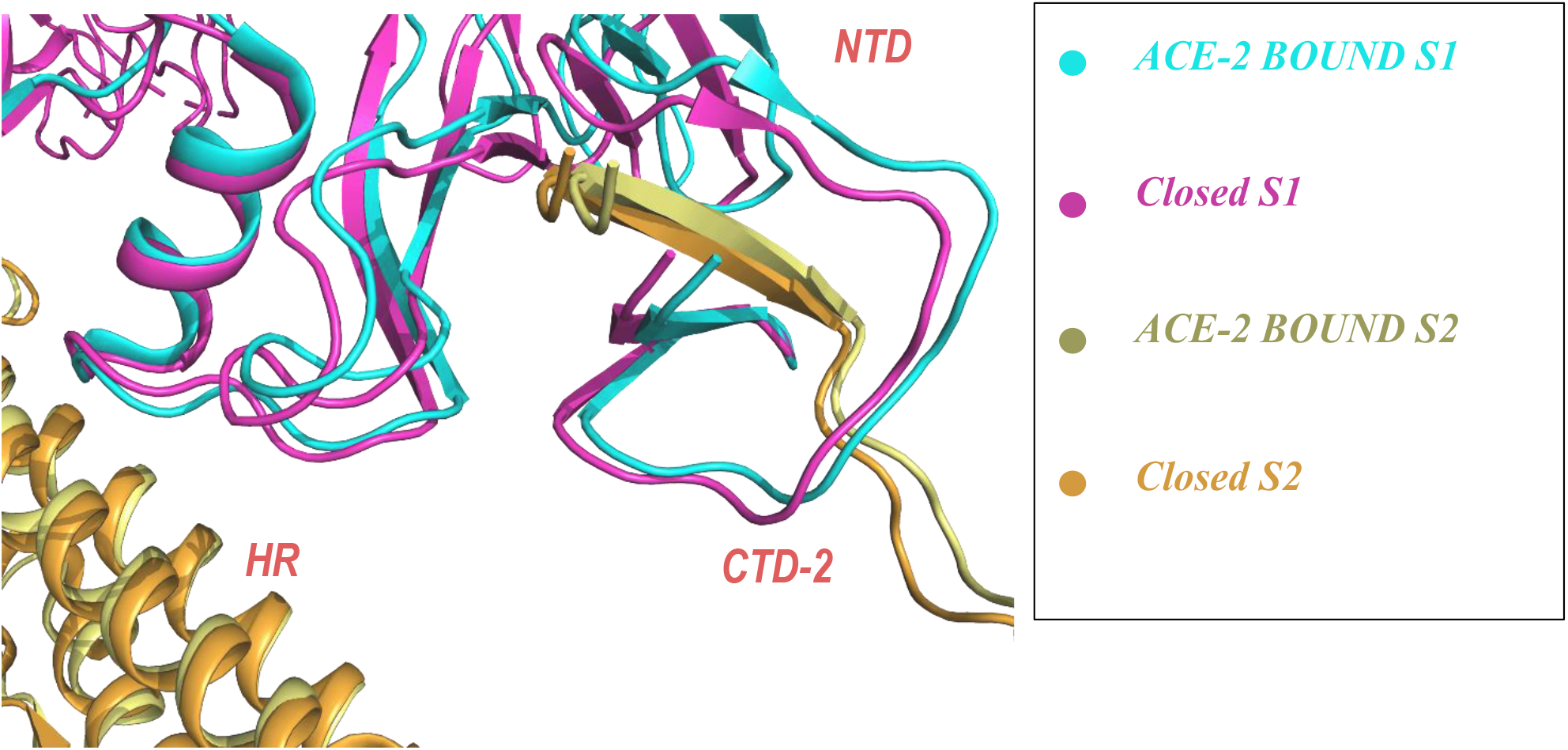
Superimposed S1/S2 interface at ‘Closed’ and ‘ACE-2 Bound’ states.

**Figure 1.9-.**
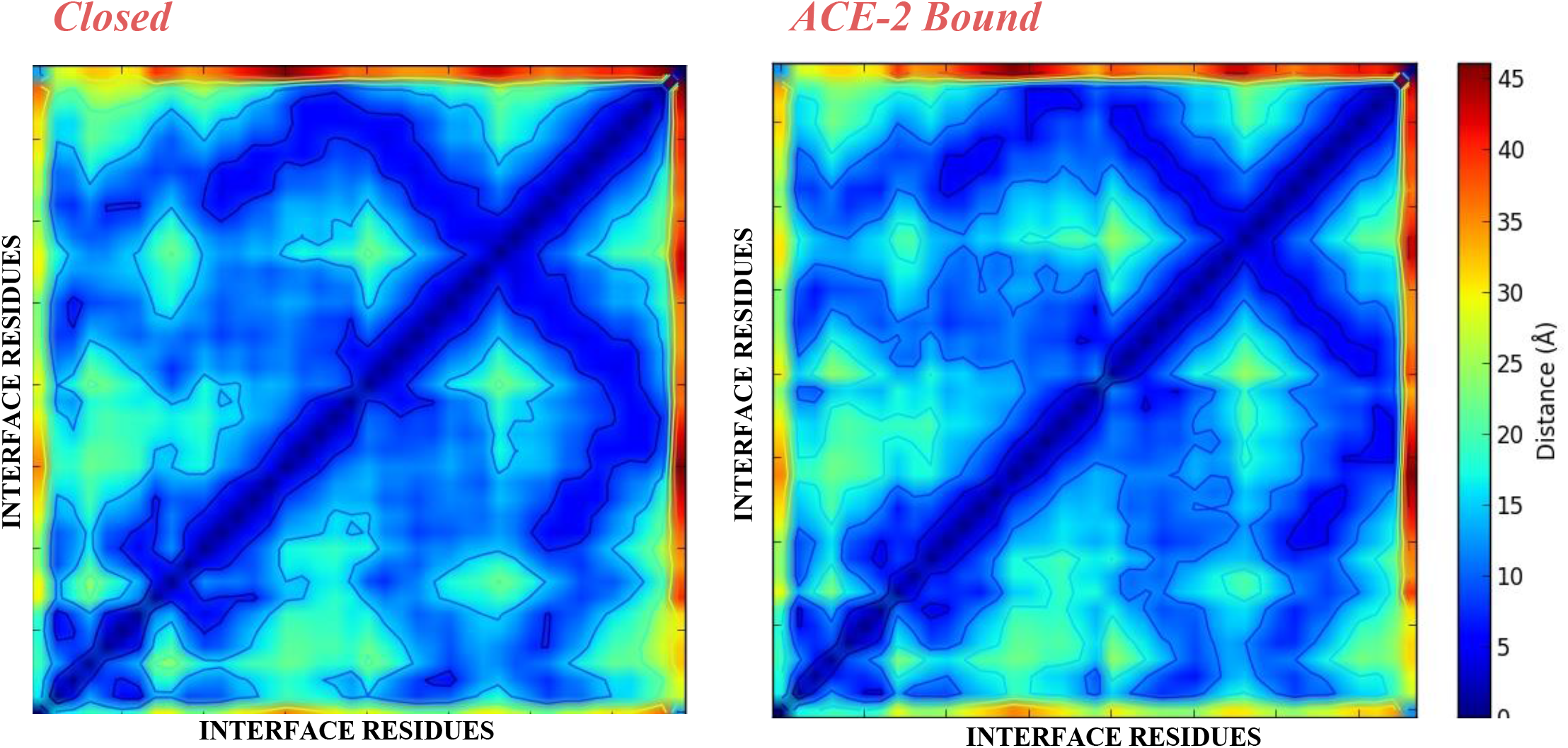
S1/S2 Distance Map confirms the fact that the relative distance between S1/S2 interface residues start diminishing in the event of ACE-2 binding.

## Materials and Methods

### i) pH and Charge Calculations

To observe pH-induced changes, we used PropKa 3.0 **[27]**. The pKa calculation was done for all the titratable residues of the different states of S protein (Chain A). Initially, we performed the calculations for the experimental S protein structures in Closed and Three RBD bound states (PDB ID: 6VXX and 6A98 respectively). To also assess the nature of S1/S2 interface of the full-length model of open and closed states, we used Amaro Lab model which is available as open-source. The pKa values of the titratable interface residues were then taken for further consideration, as our interest fits into the S1/S2 interface where possible initial events of S1 dissociation occur. Then the charge of the titratable residues upon decreasing pH *i.e*., 7 to 3 was calculated for the experimental and modelled structures to find the effect of pH in order to prove our hypothesis about unfavourable charge distribution. The charge values were obtained with the pH plot profiling module available at VMD 1.9.3. The titratable residues of S1/S2 interface were subjected for the charge profiling with the decreasing pH in the units of 0.01. The data points were then retrieved for comprehensive plotting with GNUPLOT 5.2.

### ii) Comparing Precleaved/Postcleaved States with ANM-NMA

For better understanding the roles of S1/S2 cleavage, we need to compare the S1/S2 interface environment in both the states. We chose the models which clearly depict the precleaved (CHARMM-Model)/postcleaved (Amaro Lab Model) states. For both structures, backbone Anisotropic Network Model-Normal Mode Analysis (ANM-NMA) was performed for 3 modes due to the reason that 3 modes NMA calculation is optimum for observing global fluctuations. The square fluctuations for both states were then calculated and resulting differential fluctuation values were compared. All these steps were performed with the Python-based tool-ProDy 1.10.11 **[28]**.

### iii) Distance Map Generation for the Receptor Bound/Unbound States

For deriving a conclusive proof for the receptor-induced conformational changes in the S1/S2 interface, we built S1/S2 distance maps for the experimental structures in closed as well as in Receptor bound states (PDB ID: 6VXX and 6A98 respectively). Also, generating the distance map may help us understand how receptor binding influences the S1 dissociation. With the Protein Contact Map generator **[29]**, inter-residue distances were calculated and then the final correlation map was generated.

## Conclusion

Viral glycoproteins are the enhanced fusion machinery exhibiting the diverse dynamical and structural nature of biomacromolecules. This makes them pivotal to delineate their function. The key events during viral-host cell fusion are still enigmatic and there are several unsolved questions. One such event is the dissociation of protein subunits S1 and S2 in SARS-CoV2 S protein. Understanding this spontaneous but substantial event with the aid of unconventional MD approaches which when validated by using molecular biological techniques would give us a crystal-clear perspective of how class-I fusion proteins function. Our preliminary results on key conformational changes provide an insight towards the importance of further extending our knowledge about fusion proteins and their dissociation events. Delineating fusion proteins like this can impact Structural Virology as well as promiscuous drug development.

