## Supplementary Information for "pH and Receptor Induced Conformational Changes-Implications Towards S1 Dissociation of SARS-CoV2 Spike Glycoprotein"

*i) Contact Regions of S1/S2 Interface:*

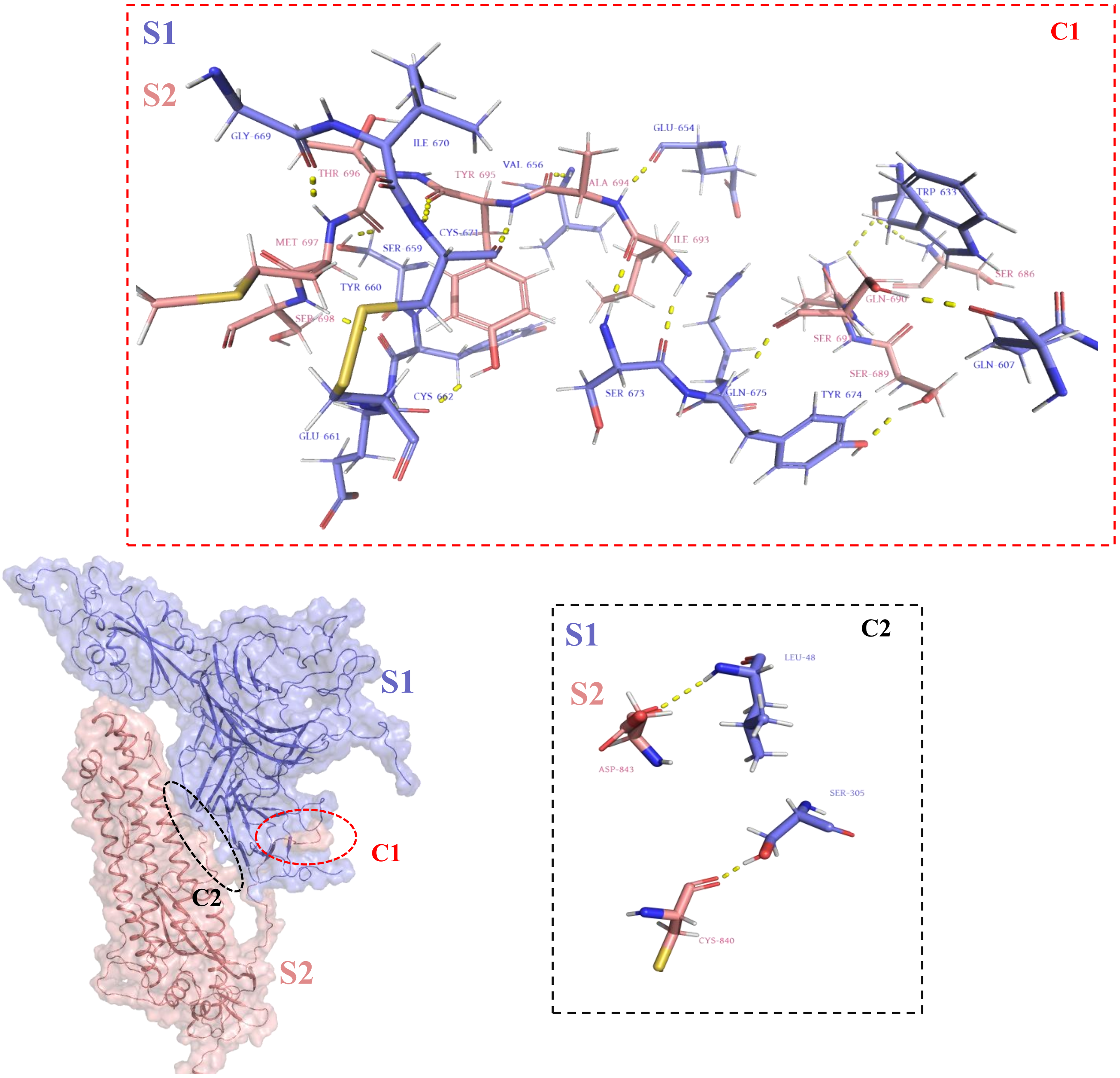


**Figure 2.1 S1/S2 interface has two contact sites – C1 with complex polar contact network and C2 with less polar contact.**

*ii) Charge Complementarity in S1/S2 Interface:*

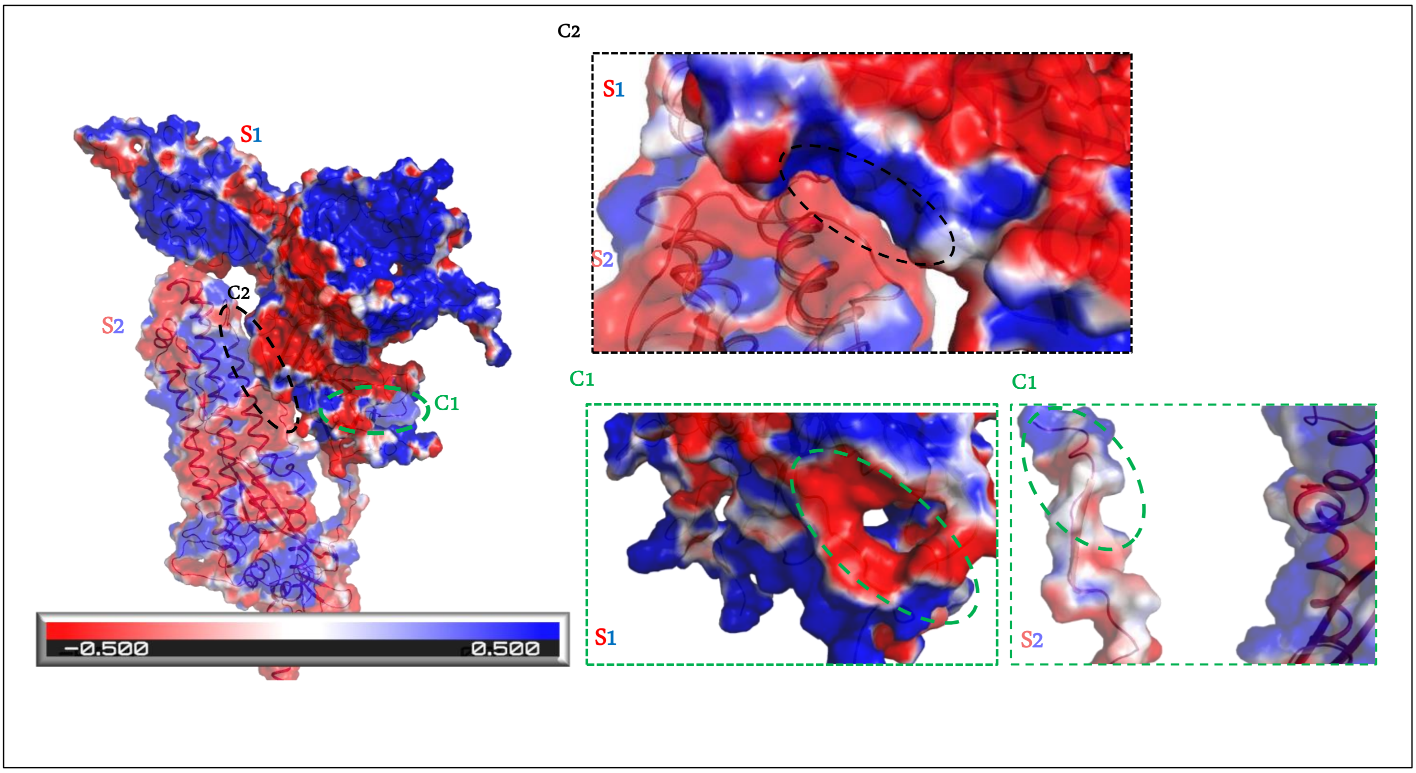


**Figure 2.2- Contact regions C1 and C2 have strong charge complementarity but they are subjected to change during the low pH**

*iii) Glycans Influencing S1/S2 Contact*

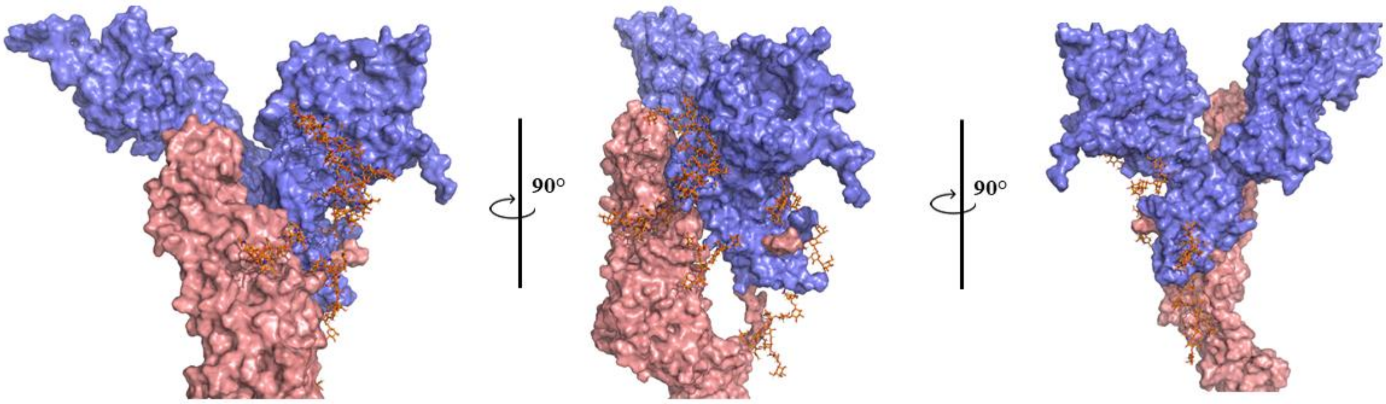


**Figure 2.3 - Glycans (shown in orange Sticks) may influence S1/S2 contact**

*iv) pKa/Charge Calculations for Amaro Lab Model Reveals the Differences Between 'Ace-2 Bound' & 'Open' States:*

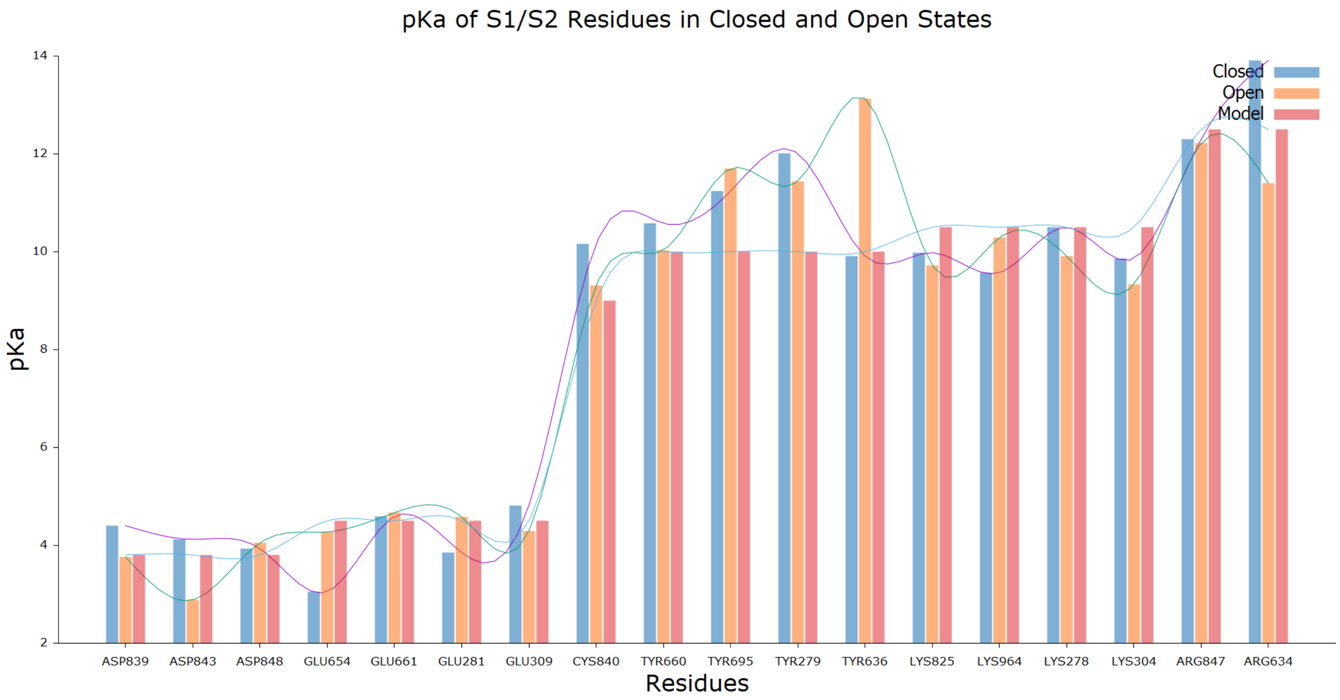


**Figure 2.4.A - pKa calculation for Amaro Lab model shows different results *i.e.*, pKa values of the interface residues of the 'Open' state are higher.**

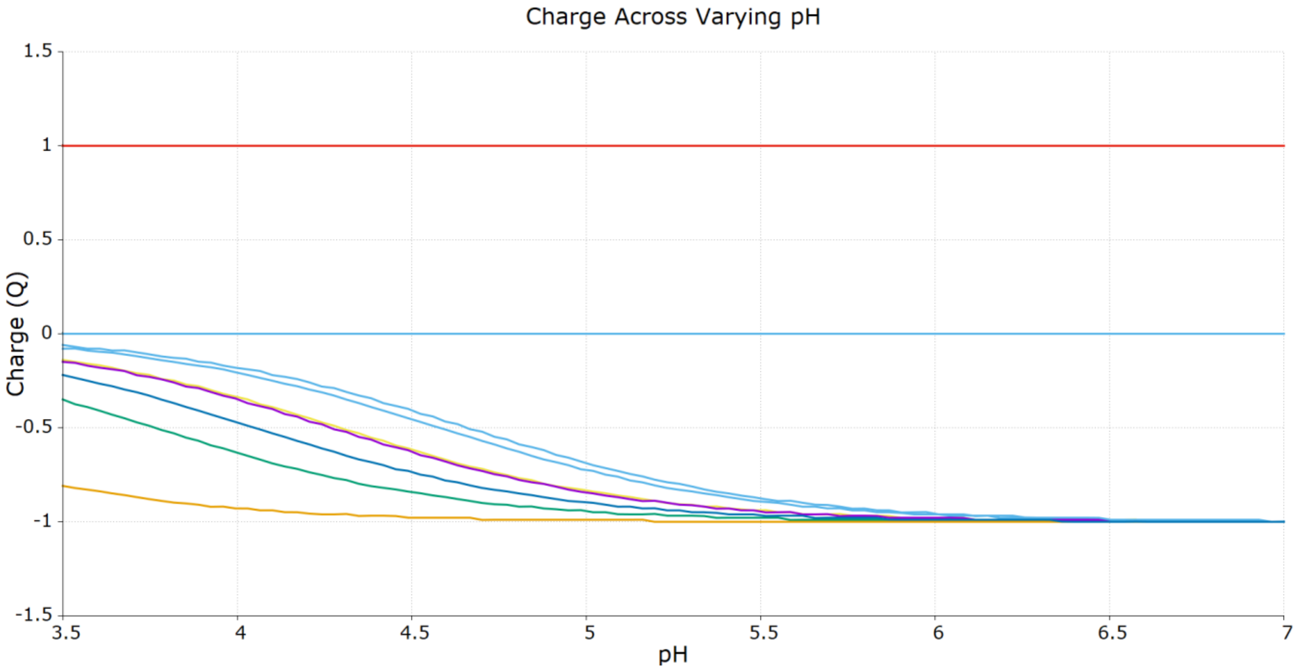


**Figure 2.4.B- Most of the interface residues (not mentioned) show increased positive charged distribution.**
